# Reconfiguration of neuronal subpopulation associated with disease-associated microglia in human Alzheimer’s disease brain

**DOI:** 10.1101/522987

**Authors:** Hongyoon Choi, Yoori Choi, Do Won Hwang, Dong Soo Lee

## Abstract

There is a growing evidence that a subset of microglia, disease-associated microglia (DAM), is crucially involved in the onset and progression of Alzheimer Disease (AD). However, it has been challenging to comprehensively know how DAM affects neuronal loss and reconstitute the complicated cellular composition. Here, we describe the relationship between neuronal reconfiguration and microglia by digitally dissecting the human brain transcriptome. We observed DAM enrichment associated with neuronal loss and the degree of amyloid deposits, which was commonly found in different brain regions. The neuronal subtype analyses showed that DAM enrichment was correlated with the proportion of two excitatory neurons located in the cortical layer 4. The direction of these relations was opposite, which implied excitatory neuronal reconfiguration in the specific cortical layer. Our results suggested immune reaction represented by DAM might mediate the reconfiguration of cellular components in AD, which may eventually have implications for diagnostics and therapeutics.

**One sentence summary:** The enrichment of disease-associated microglia signature is associated with the reconfiguration of neuronal subtypes, particularly excitatory neurons in the cortical layer 4.

## INTRODUCTION

The most common neurodegenerative disorder, Alzheimer’s disease (AD), is pathologically characterized by neuronal damage in the central nervous system (CNS) and the presence of amyloid beta deposits and neurofibrillary tangles in the brain (*1*). In spite of these characteristic pathologic markers, a major challenge is to understand the cellular and molecular mechanism of neurodegeneration and to dissect the components of CNS damage that eventually causes clinical symptoms represented by gradual memory loss. So far, the overall neuronal loss is found in the hippocampus at the early stage, which explains memory disturbance and followed by progression in the entire cerebral cortex (*2, 3*). However, as it was impossible to dissect the components of CNS damage at the cellular level, particularly for the human brain, the roles of cellular subsets in neurodegeneration were poorly understood.

Due to recent new techniques including next-generation sequencing and the single-cell transcriptome analysis, disease-associated microglia (DAM), a subset of microglia was discovered in the AD mouse model as it has a unique transcriptional signature (*4*). Further investigation also discovered the presence of DAM or DAM-like subsets of microglia in the aged human brain as well as other neurodegenerative conditions (*5, 6*). In this context, the DAM enrichment has been started to be regarded as a universal biomarker related to the neurodegeneration and aging by reflecting the increase of neurodegeneration-related molecular signals (*4, 7*). However, it is one of the great challenges to understand whether DAM plays a harmful or beneficial role in neurodegenerative disorders (*8, 9*). It is still unknown that which cellular components, particularly neurons, are affected by DAM or affecting the DAM-initiated immune reaction in the brain. In order to understand the role of DAM in CNS damages, the reconstitution of heterogeneous cellular components in the brain should be comprehensively investigated in the diseased brain. Due to the advances in various data acquisition technologies, recent studies discovered the transcriptomic atlas of the brain in the cellular level and eventually identified neuronal subtypes (*10–12*). In this regard, it is expected that we could comprehensively explore which specific neuronal subtypes, such as specific excitatory neurons, would be changed as AD progress and how this would relate to the immune reaction in the brain.

Here, we leverage the human neuronal transcriptome atlas of the brain (*10*) and tissue RNA-seq data acquired from human AD to investigate neuronal subpopulation according to the microglial landscape. The publicly available postmortem RNA-seq datasets were designed to obtain tissue transcriptome of specific brain regions, which captured the gene expression of all the constituent cells. To estimate microglial enrichment including DAM from the bulk RNA-seq data, we utilized alleged gene sets of immune cellular subsets. Furthermore, to dissect neuronal components, a digital deconvolution method and genetic profiles of neuronal subtypes were employed (*10, 13*). We aimed to investigate how the neuronal subpopulation changes with neurodegeneration and how it relates to the DAM. The comprehensive knowledge in the cellular landscape of AD brain can be a key to understand neurodegeneration and impact on the discovery of new therapeutic targets.

## RESULTS

### Disease-associated microglia associated with neuronal loss

We firstly estimated the enrichment score of disease-associated microglia (DAM) from mRNA transcriptome data of human brain tissue. Briefly, a gene set associated with human aged microglia was selected (*5*) and then the enrichment score for each sample was calculated. We also estimated the cell enrichment scores including neurons, overall macrophage, M1-like and M2-like cells using predefined gene sets (*14*). The data were obtained from a cohort of AD and control brains from the Mount Sinai Brain Bank (MSBB) which included four brain regions, parahippocampal gyrus (PHG), prefrontal cortex (PFC), superior temporal gyrus (STG) and inferior frontal gyrus (IFG), and Mayo Clinic Brain Bank RNA-seq study obtained from the temporal cortex. Note that the following analyses were mainly performed in PHG where the neuronal loss was found at the early stage (*2, 15*), and results for other brain regions were also compared. As a result, the DAM enrichment score was closely related to AD brain (**Fig. 1A**). The DAM enrichment scores of AD was significantly higher than those of controls in PHG (**Fig. 1B**) (0.26 ± 0.98 vs −0.61 ± 0.76 for AD and control, respectively; t = 7.20, p < 1×10^−13^). The higher DAM enrichment was also found in other brain regions including PFC, STG and IFG (**Fig. S1**). The enrichment scores of DAM and neuron were negatively correlated to each other regardless of brain regions (r = −0.77, p < 1×10^−15^ for PHG, r = −0.63, p < 1×10^−15^ for PFC, r= −0.71, p < 1×10^−15^ for STG, r = −0.70, p < 1×10^−15^ for IFG and r= −0.61, p < 1×10^−14^ for Mayo-temporal) (**Fig. 1C, Fig. S2**). Amyloid plaque deposits were positively associated with DAM enrichment score (r = 0.41, p < 1×10^−9^ for PHG, r = 0.26, p < 1×10^−4^ for PFC, r = 0.33, p < 1×10^−7^ for STG, and r = 0.17, p < 0.01 for IFG) (**Fig. 1D, Fig. S3**).

**Figure 1.**
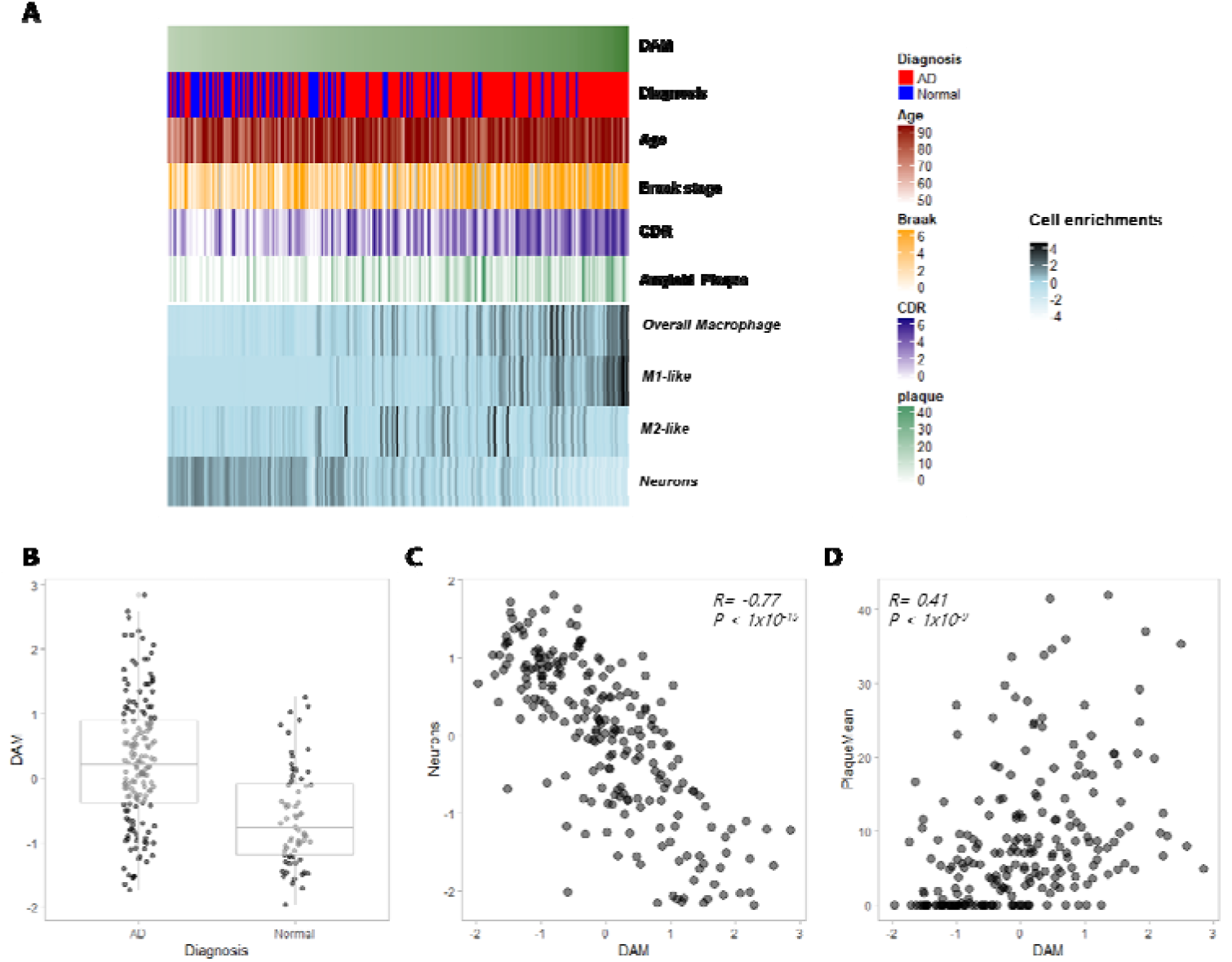
Disease associated microglia (DAM) enrichment score associated with Alzheimer’s disease (AD) pathology in human brain. (A) DAM enrichment score was calculated by a specific gene set of aged microglia and tissue RNA-sequencing data acquired from parahippocampal gyrus (PHG). We also estimated immune cellular signatures including macrophage, M1- and M2-like cells as well as the neuron signature and then plotted according to the order of DAM enrichment. (B) The DAM enrichment scores of AD brain was significantly higher than those of controls in PHG (0.26 ± 0.98 vs −0.61 ± 0.76 for AD and control, respectively; t = 7.20, p < 1×10^−13^). (C) The correlation between the neuron and DAM signatures, which showed a significant negative correlation (r = −0.77, p < 1× 10^−15^). (D) The degree of amyloid plaques was positively associated with DAM enrichment score (r = 0.41, p < 1×10^−9^).

### DAM and macrophage-related signatures

As DAM is a subset of overall microglia in the brain, we estimated the relationship between brain immune cell signatures, DAM and other macrophage-related signatures. The macrophage-related signatures were estimated by specific gene sets of cellular markers (The cellular markers are summarized in **Table S1**) (*14*). The DAM enrichment score was positively correlated with overall macrophage and M1-like cell population in PHG (r = 0.63 and r = 0.74 for macrophage-like and M1-like cells, respectively, **Fig. 2A**). It was not correlated with M2-like cell population. The data of AD patients were selected and divided into 4 subgroups according to the immune cell subpopulation of M1-like and M2-like cells: M1^+^M2^+^, M1^+^M2^-^, M1^-^M2^+^, and M1^-^M2^-^ types. ‘Positive’ and ‘negative’ represented the relatively more enriched M1-like or M2-like cells than others and were determined by the median value of each enrichment score. There was a significant effect of the subgroups on DAM and neuronal loss (F(3, 154) = 54.2, p < 1×10^−15^ for DAM and F(3, 154) = 26.9, p < 1×10^−13^ for neurons). The DAM enrichment score was the highest in the M1^+^M2^-^ subgroup and the enrichment score of neurons was also the lowest in the M1^+^M2^-^ subgroup (Posthoc Tukey, p < 0.05 for all comparison) (**Fig. 2B, C**). The pattern, M1^+^M2^-^ subgroup with the highest neuronal loss and DAM, was consistently found in the independent data cohort (Mayo temporal cortex) as well as other brain regions (PFL, STG and IFG) (**Fig. S4**). We plotted M1-like and M2-like subpopulation according to DAM enrichment scores and fitted by local polynomial regression (Loess fitting) (**Fig. 2E**). According to the progression of DAM enrichment, both M1-like and M2-like signatures were initially increased, while further increased DAM was associated with decreased M2-like cells and increased M1-like cells in all brain regions (**Fig. S5**). M2-like cells were decreased according to DAM for Mayo temporal cortex data (**Fig. S5**).

**Figure 2.**
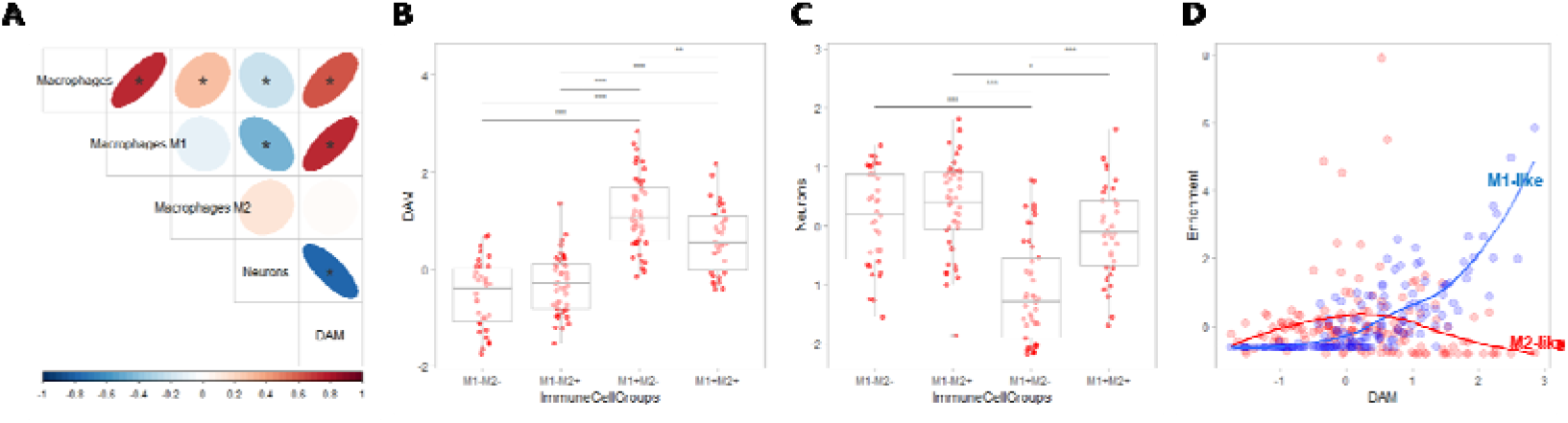
The relationship between immune cell signatures estimated by tissue RNA-sequencing. (A) The DAM enrichment score was compared with other immune cell signatures estimated by predefined gene signatures of macrophage, M1 and M2 cells. The DAM enrichment was positively associated with macrophage and M1 enrichment, while not associated with M2-like cell. (B, C) Using the median value of M1- and M2-like cells enrichment, samples were divided into 4 subgroups: M1^+^M2^+^, M1^+^M2^-^, M1^-^M2^+^, and M1^-^ M2^-^. The M1^+^M2^-^ subgroup showed the highest DAM enrichment score (B) and the lowest neuron signature, which implied the highest neuronal loss (C). M1-like and M2-like scores were plotted with DAM enrichment score and fitted by local polynomial regression (D). According to DAM enrichment, both scores were initially increased and then the M2-like cell score was decreased while the M1-like cell score was still increased.

### Reconfiguration of neuronal subpopulation associated with DAM

As overall neuronal loss was closely associated with DAM, we investigated whether DAM affected a composition of neuronal subtypes. Using gene profiles of excitatory and inhibitory neurons of the human brain cortex (*10*), we calculated the relative population of neuronal subtypes for each sample (**Fig. 3**). The gene profile for each neuronal subtype is summarized in **Table S2**. The subpopulation was represented by a heatmap (**Fig. 3A**) and barplots for PHG and other brain regions (**Fig. 3B, Fig. S6**). Neuronal subtypes which occupied 0% in most samples were excluded from the further analysis. We calculated the correlation between DAM enrichment and each neuronal subtypes. In PHG, DAM was negatively correlated with Ex2 neurons and positively correlated with Ex3 neurons (r = −0.55, p < 1×10^−15^ for Ex2 and r = 0.55, p < 1×10^−15^ for Ex3) (**Fig. 4A**). Notably, Ex2 and Ex3 neurons constitute the cortical layer 4 (*10*), thus, this opposite direction of correlation meant repopulation of cortical layer 4 associated with DAM enrichment. The positive correlation of Ex3 and negative correlation of Ex2 to DAM were commonly found in other brain regions (**Fig. S7**). We also compared the neuronal subpopulation between AD and control brains. A volcano plot represented increased and decreased neuronal subpopulation in AD compared with control brain (**Fig. 4B**). The proportion of Ex3 neuron was significantly increased and the proportion of Ex2 neuron was significantly decreased in AD (Ex2: 0.10 vs. 0.15, t = −4.3, p < 0.0001; Ex3: 0.06 vs. 0.03, t = 5.0, p < 1×10^−5^ for AD and control, respectively) (**Fig. 4C and 4D**). Increased Ex3 and decreased Ex2 neuronal proportion in AD were also found in PFL, while this finding was not definite in other brain regions including STG and IFG (**Fig. S8**). Of note, the proportions of Ex2 and Ex3 were associated with DAM in all brain regions including STG and IFG.

**Figure 3.**
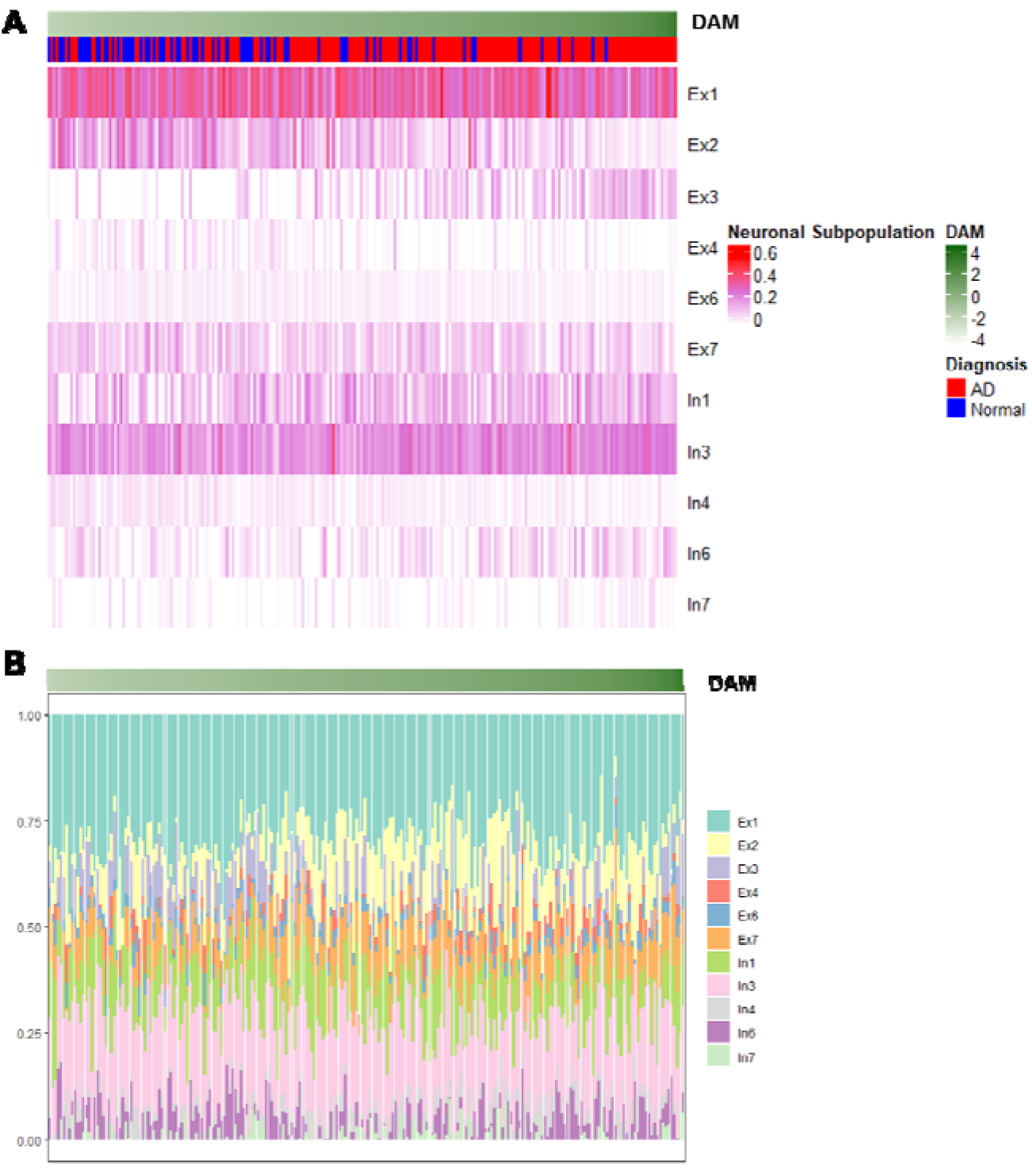
Reconfiguration of neuronal subpopulation associated with DAM enrichment. Using genetic profiles of 8 excitatory and 8 inhibitory neurons, proportional changes of the subpopulation in the brain (PHG) in accordance with DAM enrichment were estimated. A heatmap (A) and a barplot (B) showed relatively increased Ex3 neuronal proportion and decreased Ex2 neuronal proportion according to DAM enrichment score.

**Figure 4.**
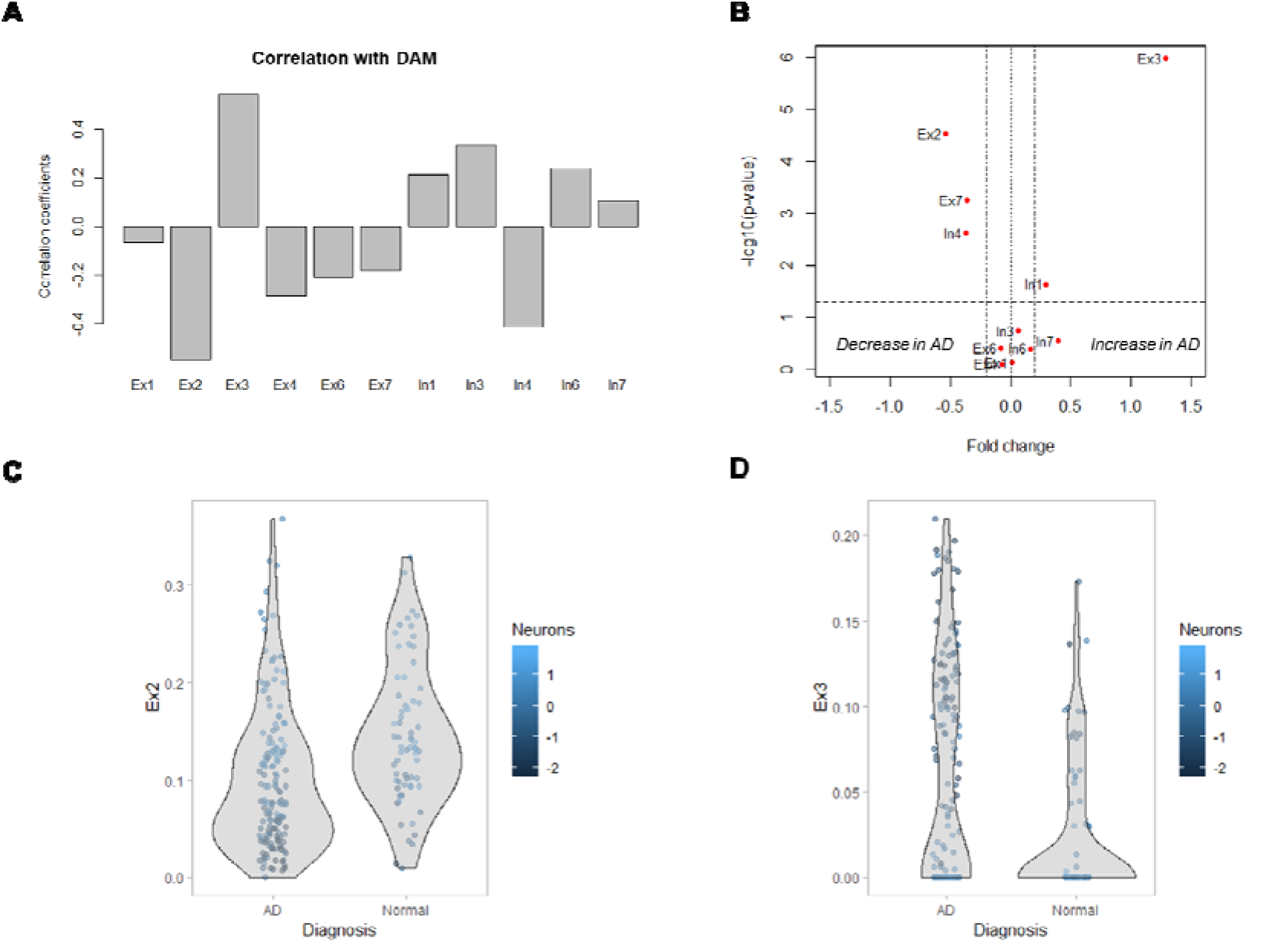
Proportional changes of two excitatory neurons associated with DAM. (A) Two excitatory neurons in the cortical layer 4, Ex2 and Ex3, were correlated with DAM enrichment score. DAM was negatively correlated with Ex2 neurons and positively correlated with Ex3 neurons in the PHG (r = −0.55 and r = 0.55, respectively; p < 1×10^−15^). (B) The proportions of neuronal subtypes of AD and normal controls were compared and plotted with a volcano plot. (C, D) The proportion of Ex2 neuron was decreased and that of Ex3 neuron was increased in AD brain. The proportion of Ex3 neuron was significantly increased and the proportion of Ex2 neuron was significantly decreased in AD (Proportion of Ex2: 0.10 vs. 0.15, t = −4.3, p < 1×10^−4^; Proportion of Ex3: 0.06 vs. 0.03, t = 5.0, p < 1×10^−5^ for AD and control, respectively).

## DISCUSSION

In this study, we found the reconfiguration of cellular subsets, particularly neurons, in the human AD brain according to the DAM enrichment. The overall neuronal loss was closely associated with the DAM regardless of the brain region. The signatures of immune cell subsets including DAM, M1-like and M2-like cells were compared. We found the different patterns of immune cell enrichment according to the AD progression. The neuronal subpopulation was estimated by digitally dissecting the neuronal components. Among 8 excitatory and 8 inhibitory neuronal subtypes, the proportions of two excitatory neurons, Ex2 and Ex3, located on the cortical layer 4 were significantly associated with DAM enrichment. Increased DAM was positively correlated with the proportion of Ex3 while it was negatively correlated with the proportion of Ex2, which suggested the excitatory neuronal reconfiguration in the cortical layer 4. The result of the cellular landscape suggested the close link of microglia-mediated immune reaction and neuronal reconfiguration as well as overall neuronal loss in the AD brain.

The DAM enrichment was closely associated with overall neuronal damage and amyloid plaques in the human brain. This finding was consistent with the previous finding which suggested the synaptic loss in the AD mouse model according to the microglial activation and complement-mediated immune reaction (*16*). This microglia-mediated neuronal loss has been one of the therapeutic targets as microglial depletion protected neuronal loss and cognitive symptoms in the animal models (*17, 18*). Furthermore, the association between DAM and amyloid plaques corresponds to the animal pathologic study, which showed the co-localized DAM and amyloid plaques (*4*). In spite of the positive correlation between DAM and amyloid plaques, there is still controversy in the role of DAM in AD pathology regarding neuronal protection or neuronal damage (*19*). Amyloid plaques can lead to microglial phagocytic activation for the protection or induce neuronal damage mediated by a detrimental immune reaction. Although our cross-sectional analysis hardly explains the exact role of DAM in amyloid deposition, the close association of DAM, amyloid plaques and neuronal loss in the human brain implies the cascades of this AD-related immune reaction and neuronal loss, which could eventually lead to clinical symptoms. Moreover, the neuronal reconfiguration associated with DAM in our results could support the idea of microglia-mediated neuronal damage (*20, 21*).

Following the close relationship between DAM and neuronal cell loss, the next key question at the cellular level is how AD pathology reconfigures the neuronal subpopulation. Our finding suggests the reconstitution of excitatory neurons in the specific layer, cortical layer 4. DAM enrichment was closely related to the neuronal proportion of Ex2 and Ex3, but the direction was opposite. According to the progression of DAM enrichment, the Ex3 neuronal proportion was increased while Ex2 neuronal proportion was decreased. Of note, the difference in gene expression profiles of two excitatory neurons was associated with the connectivity (*10*). The connection abnormality of excitatory neurons caused by the reconfiguration of excitatory neurons in layer 4 could be related to the excitatory imbalance in the microcircuit of AD brain (*22, 23*). More specifically, glutamate-mediated calcium overload induced by amyloid plaques result in the excitatory imbalance and eventually glutamate toxicity (*24*). Of note, relatively increased Ex3 proportion and decreased Ex2 proportion in AD brain were found in the PHG and PFL, but not in the STG, IFG and the neocortex of the temporal lobe. These two brain regions, PHG and PFL, are vulnerable brain regions according to the neurodegeneration, which leads to abnormal energy metabolism (*25*). Considering the correlation of Ex2 and Ex3 proportion to DAM regardless of the brain regions, the abnormal connectivity of excitatory neurons in the layer 4 associated with DAM would occur in the entire brain. However, due to the difference of vulnerability in neuronal damage, both PHG and PFL might have a clear association with the diagnosis of the AD. Moreover, these brain regions are related to the low efficiency of energy metabolism compared to brain connection, thus, the connections could be vulnerable to pathologic stress such as amyloid deposits (*26, 27*). Our hypothesis of excitatory imbalance and abnormal connectivity in the cortical layer 4 according to DAM should be clarified in further *in vitro* and animal studies. Nonetheless, our results suggest a plausible explanation for the dynamic neuronal reconfiguration due to the immune reaction in the brain. Furthermore, it could support the idea of the regional variability in the neuronal damage associated with excitatory imbalance and metabolic demands, and complicated neuronal connectivity disruption of the AD brain (*28*).

The comparison of immune cell signatures in the brain suggested the changes of microglial subtypes according to the disease progression. Two different macrophage signatures, M1-like and M2-like cellular enrichment scores were differently associated with DAM and neuronal loss patterns. More specifically, the M1-like cell enrichment score was closely associated with DAM and neuronal loss, while M2-like cell enrichment score was increased in samples with relatively low DAM enrichment but then decreased according to DAM enrichment. Thus, when we divided the samples by the enrichment scores of M1-like and M2-like cells, the neuronal loss and DAM were the highest in the M1^+^M2^-^ subgroup. This finding corresponds to the finding of three microglial subtypes in AD mouse model, which showed homeostatic microglia, stage 1 and stage 2 DAM (*4*). The report suggested that the composition of three microglial subtypes were dynamically changed according to the disease progression (*4*). In this context, initially increased and then decreased patterns of M2-like cell enrichment score in our results could reflect the dynamic transitional state of microglia in the diseased brain (*29*). Further human microglial subtypes using single cell analyses in AD brain might elucidate the dynamic state of microglia and its role in AD progression. As a single cell-level atlas of human microglia could provide genomic profiles of various state of microglia, another future work of digitally dissecting microglial subtypes in tissue RNA-seq data could further clarify the cellular landscape of neurodegeneration.

We investigated the association of DAM enrichment and neuronal reconfiguration in the human brain. Since it is difficult to analyze a number of human brain samples with the different disease state in cellular level, we leveraged immune cell signatures and gene expression profiles of neuronal subtypes previously dissected using single cell analyses. As a result, we found the close relationship between DAM enrichment and overall neuronal loss. By comparing immune cellular signatures, we suggested the changes of microglial subtypes enrichment according to the disease progression. More importantly, we suggested that the DAM enrichment was associated with the reconfiguration of excitatory neuronal subtypes in the specific cortical layer, which supports the idea of abnormal connectivity mediated by an excitatory imbalance in AD pathophysiology. The result of the close relationship of microglia-mediated immune reaction, neuronal loss, and cellular reconfiguration could provide an insight into how AD-related pathologic features including amyloid plaques affect clinical symptoms related to specific neuronal damages.

## MATERIALS AND METHODS

### Subjects and samples

The Mount Sinai Brain Bank (MSBB) RNA-seq study was used for the analysis (synapse ID: syn3157743 from the Accelerating Medicines Partnership - Alzheimer’s Disease Knowledge Portal) (*30*). Post-mortem brain samples were collected from the four different brain regions, PHG (Brodmann Area 36), PFC (Brodmann Area 10), STG (Brodmann Area 22), and IFG (Brodmann Area 44). Tissue RNA-seq was generated using the TruSeq RNA Sample Preparation Kit v2 and Ribo-Zero rRNA removal kit (Illumina, San Diego, CA). For RNA-seq data processing, raw sequence reads were aligned to human genome reference hg19 with star aligner (v2.3.0e) and reads were summarized to gene level counts using the featureCounts (*31*).

Mayo Clinic Brain Bank RNA-seq study was also used for additional data (synapse ID: syn5550404 from the Accelerating Medicines Partnership - Alzheimer’s Disease Knowledge Portal). RNA-seq based transcriptome data were generated from post-mortem temporal cortex samples of AD patients, controls, patients with pathologic aging and progressive supranuclear palsy patients. We selected RNA-seq data of AD and controls (n = 82 and 78, respectively). Detailed process pipelines are described in the previous report (*32*).

### Disease-associated microglia enrichment

To define disease-associated microglia (DAM) in the human brain, we used a gene set that is preferentially expressed by microglia in the aged human brain, which also associated with pathological processes (*5, 7*). To obtain DAM enrichment score, we used a single sample gene set enrichment analysis (ssGSEA) which provide the activity of DAM signature for each sample (*33*). DAM signature was provided by the previous study (*5*), by selecting 650 genes which highly expressed in human aged microglia cells with a 4-fold increase and adjusted p < 0.05. The output of ssGSEA was normalized by z-score across samples and used as a DAM enrichment score.

### Brain immune cells and neurons enrichment

To evaluate the subpopulation of immune cells in the brain tissue, cell types enrichment scores were evaluated. A gene-signature based method for inferring cell types from tissue transcriptome profiles, xCell tool (http://xcell.ucsf.edu/), was used (*34*). xCell infers more than 60 cell types, however, most cell types are not associated with brain tissue. Thus, we selected macrophage-related signatures, whole macrophage, M1 and M2 signatures, and neurons signatures. The gene expression data of brain tissue samples were used to estimate macrophage-like, M1-like and M2-like cell enrichment scores.

### Clusters based on immune cell subtype

Immune cell enrichment scores of the brain tissue of AD patients were selected. To find different cell population features including DAM and neuronal loss, samples were clustered based on the brain immune cell subtypes, M1-like and M2-like cells. Relatively M1-like cell enriched groups were defined by the median value of the enrichment score. By using the two respective threshold values of M1-like and M2-like cell enrichment scores, 4 subgroups were defined: M1+M2^+^, M1^+^M2^-^, M1^-^M2^+^, and M1^-^M2^-^ types. ‘Positive’ and ‘negative’ represents the sample with higher and lower than the median value of enrichment score, respectively.

### Neuronal subtype signatures

Neuronal subtype signatures were obtained by the single cell RNA-seq data of human brain neurons (*10*). The data consist of 3227 sets of single neuron RNA-seq data acquired by isolated cellular nuclei. Accordingly, 16 neuronal subtypes, 8 excitatory and 8 inhibitory neurons, were defined by using clustering analysis. Detailed methods are described in the previous report. Briefly, for RNA-seq data processing, short reads were aligned to human hg38 reference genome (STAR ver 2.3.0) and quantified by HTSeq (ver 0.5.4p5). Transcription levels were then converted to transcript per million mapped reads (TPM). Gene expression signatures of each neuron subtype were estimated by median TPM of all marker genes. These representative gene expression data of 16 neuronal subtypes were used as reference gene expression data for digitally dissecting cellular subtypes from bulk RNA-seq data.

### Digital dissection of neuronal subtypes from tissue samples

To dissect the neuronal subtypes from bulk RNA-seq data acquired from postmortem human brain, CIBERSORT was used (*13*). CIBERSORT deconvolute gene expression data of tissues into estimated subpopulation of 16 neuronal subtypes using aforementioned single cell-level transcriptomic data. The tissue RNA-seq data were resampled to have same gene identifiers with the reference gene signatures of 16 neuronal subtypes and these rearranged data were supplied to CIBERSORT as inputs. A subpopulation of 16 neuronal subtypes was identified by default parameters of the algorithm.

### Statistical analysis

The correlation between two continuous variables was estimated by Pearson’s correlation. The comparison of DAM enrichment scores between two diagnostic groups (AD and controls) was conducted by the two-sample t-test. 16 neuronal subtypes population of AD and controls were also compared using the two-sample t-test. To compare the statistical results of 16 neuronal subtypes, volcano plots were drawn by plotting fold changes and p-values calculated by the two-sample t-tests between AD and controls. The comparison of DAM and neuron enrichment scores between four immune cell-based clusters was conducted by one-way ANOVA, followed by the post hoc Tukey’s test.

## SUPPLEMENTARY MATERIALS

Figure S1. DAM enrichment scores of AD and controls in various brain regions.

Figure S2. DAM enrichment scores associated with neuronal loss in various brain regions.

Figure S3. DAM enrichment scores associated with amyloid plaques in various brain regions.

Figure S4. Subgroups divided by M1- and M2-like cell enrichment.

Figure S5. M1- and M2-like cell signatures according to DAM enrichment in various brain regions.

Figure S6. The proportion of neuronal subpopulation according to DAM enrichment in various brain regions.

Figure S7. Correlation coefficients between the proportion of neuronal subpopulation and DAM enrichment.

Figure S8. Volcano plots for the comparison of the proportion of neuronal subtypes of AD and controls.

Table S1. Gene lists of immune cellular markers

Table S2. Gene signatures of 16 neuronal subtypes

## ACKNOWLEDGEMENT

The results here are in whole based upon data publicly available generated by Mount Sinai Brain Bank (MSBB) and Mayo Clinic Brain Bank.

MSBB data were generated from postmortem brain tissue collected through the Mount Sinai VA Medical Center Brain Bank and were provided by Dr. Eric Schadt from Mount Sinai School of Medicine.

Mayo Clinic Brain Bank data were provided by the following sources: The Mayo Clinic Alzheimers Disease Genetic Studies, led by Dr. Nilufer Ertekin-Taner and Dr. Steven G. Younkin, Mayo Clinic, Jacksonville, FL using samples from the Mayo Clinic Study of Aging, the Mayo Clinic Alzheimers Disease Research Center, and the Mayo Clinic Brain Bank. Data collection was supported through funding by NIA grants P50 AG016574, R01 AG032990, U01 AG046139, R01 AG018023, U01 AG006576, U01 AG006786, R01 AG025711, R01 AG017216, R01 AG003949, NINDS grant R01 NS080820, CurePSP Foundation, and support from Mayo Foundation. Study data includes samples collected through the Sun Health Research Institute Brain and Body Donation Program of Sun City, Arizona. The Brain and Body Donation Program is supported by the National Institute of Neurological Disorders and Stroke (U24 NS072026 National Brain and Tissue Resource for Parkinsons Disease and Related Disorders), the National Institute on Aging (P30 AG19610 Arizona Alzheimers Disease Core Center), the Arizona Department of Health Services (contract 211002, Arizona Alzheimers Research Center), the Arizona Biomedical Research Commission (contracts 4001, 0011, 05-901 and 1001 to the Arizona Parkinson’s Disease Consortium) and the Michael J. Fox Foundation for Parkinsons Research.

## FUNDING

This research was supported by the National Research Foundation of Korea (NRF) grant funded by the Korean Government (MSIP) (No. 2017M3C7A1048079). This research was also supported by a grant of the Korea Health Technology R&D Project through the Korea Health Industry Development Institute (KHIDI), funded by the Ministry of Health & Welfare, Republic of Korea (HI14C0466), and funded by the Ministry of Health & Welfare, Republic of Korea (HI14C3344), and funded by the Ministry of Health & Welfare, Republic of Korea (HI14C1277), and the Technology Innovation Program (10052749).

## AUTHOR CONTRIBUTIONS

H.C., D.S.L. designed the study. H.C. performed the computational analysis. Y.C. and D.W.H. contributed the data analyses. H.C. and D.S.L. wrote the paper. All authors read and approved the final manuscript.

## COMPETING INTERESTS

The authors declare no competing financial interests.

## DATA and MATERIALS AVAILABILITY

The Mount Sinai Brain Bank (MSBB) and Mayo Clinic Brain Bank RNA-seq data can be downloaded from Accelerating Medicines Partnership - Alzheimer’s Disease Knowledge Portal (synapse ID: syn3157743 and syn5550404).

The single cell-level human brain transcriptome atlas data can be downloaded from dbGAP (https://www.ncbi.nlm.nih.gov/projects/gap) with accession number phs000833.v3.p1. Processed data can be downloaded from http://genome-tech.ucsd.edu/public/Lake_Science_2016/.

