## Supplementary Figures for "Reconfiguration of neuronal subpopulation associated with disease-associated microglia in human Alzheimer’s disease brain"

**Additional File**

### Supplementary Figures

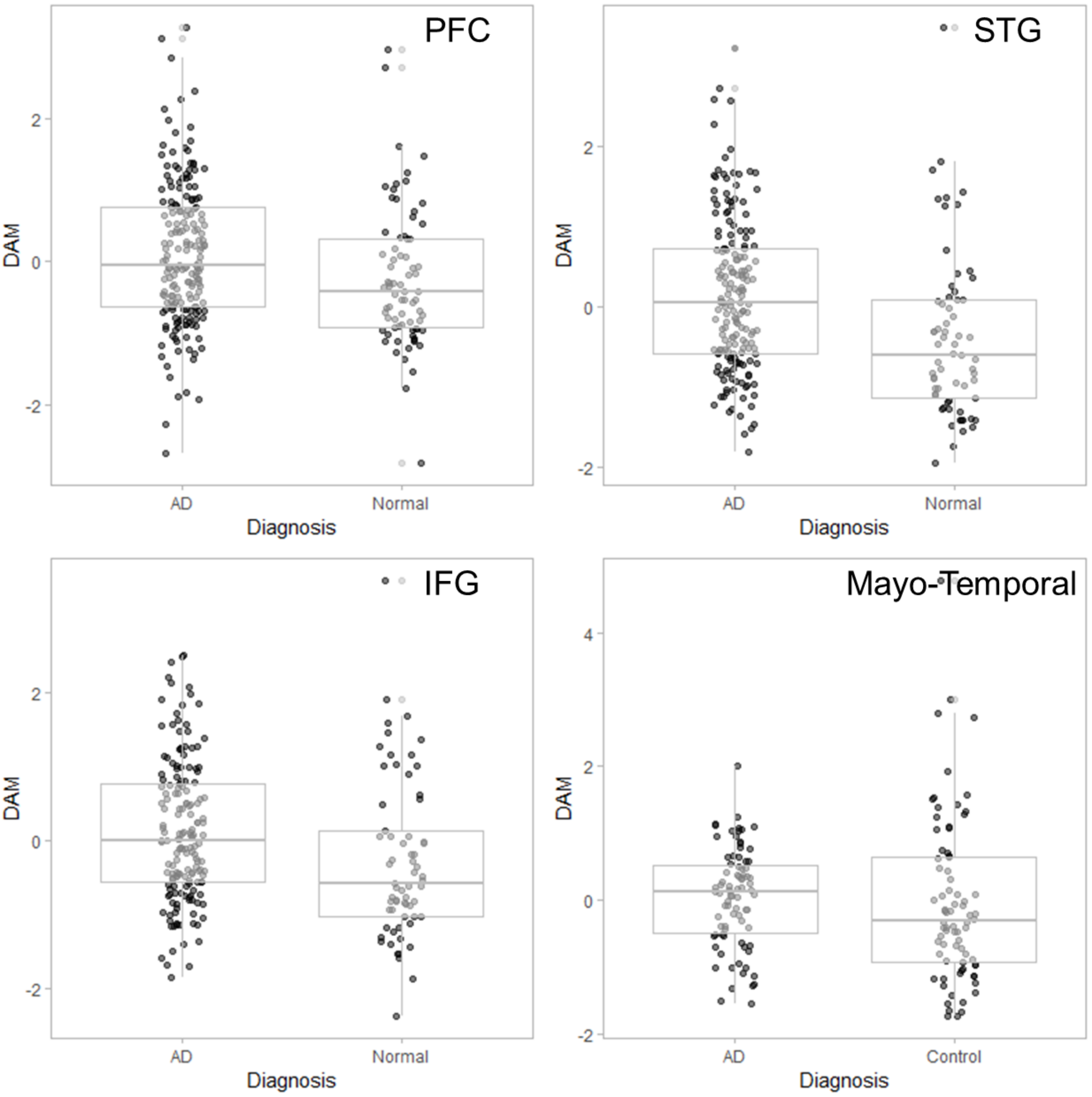


**Figure S1. DAM enrichment scores of AD and controls in various brain regions.** DAM enrichment scores of AD were significantly higher than those of controls in PFC (0.10 ± 1.00 vs -0.25 ± 0.97 for AD and control; t = 2.58, p = 0.01), STG (0.15 ± 0.95 vs -0.39 ± 1.02; t= 3.71, p = 0.0003), and IFG (0.11 ± 0.95 vs -0.28 ± 1.08; t= 2.55, p =0.01). The higher DAM enrichment score in AD than control was also found in temporal cortex of Mayo postmortem data, though it did not reach statistical significance (0.04 ± 0.73 vs -0.05 ± 1.22; t= 0.55, p = n.s.).

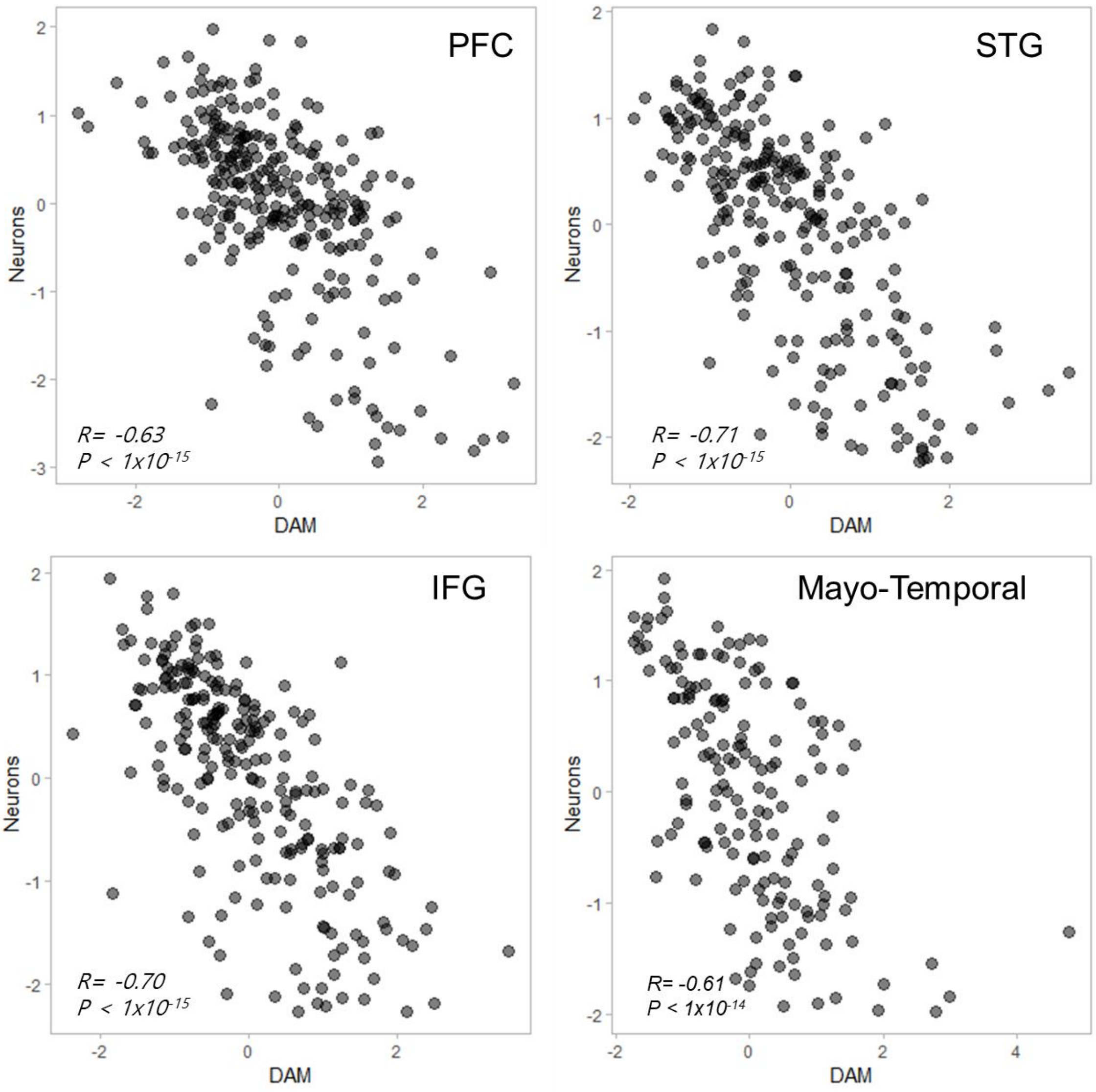


**Figure S2. DAM enrichment scores associated with neuronal loss in various brain regions.** Overall neuronal enrichment was estimated and correlated with DAM enrichment scores. The neuronal enrichment score was significantly negatively correlated with DAM score in all brain regions ($r = -0.63$, $p < 1\times10^{-15}$ for PFC, $r= -0.71$, $p < 1\times10^{-15}$ for STG, $r = -0.70$, $p < 1\times10^{-15}$ for IFG and $r= -0.61$, $p < 1\times10^{-14}$ for Mayo-temporal).

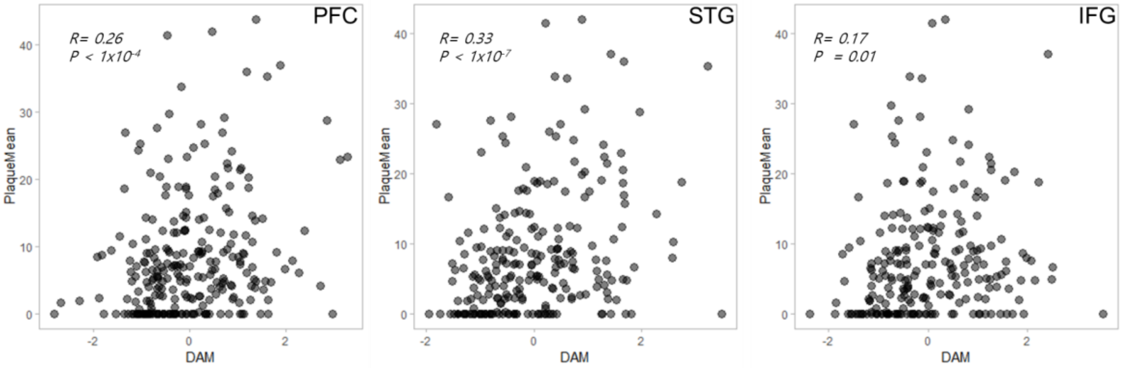


**Figure S3. DAM enrichment scores associated with amyloid plaques in various brain regions.** The density of amyloid plaques estimated by mass spectrometry was correlated with DAM enrichment score. Amyloid plaque deposits were positively correlated with DAM enrichment score in all brain regions of MSBB data ($r = 0.26$, $p < 1\times10^{-4}$ for PFC, $r = 0.33$, $p < 1\times10^{-7}$ for STG, and $r = 0.17$, $p < 0.01$ for IFG).

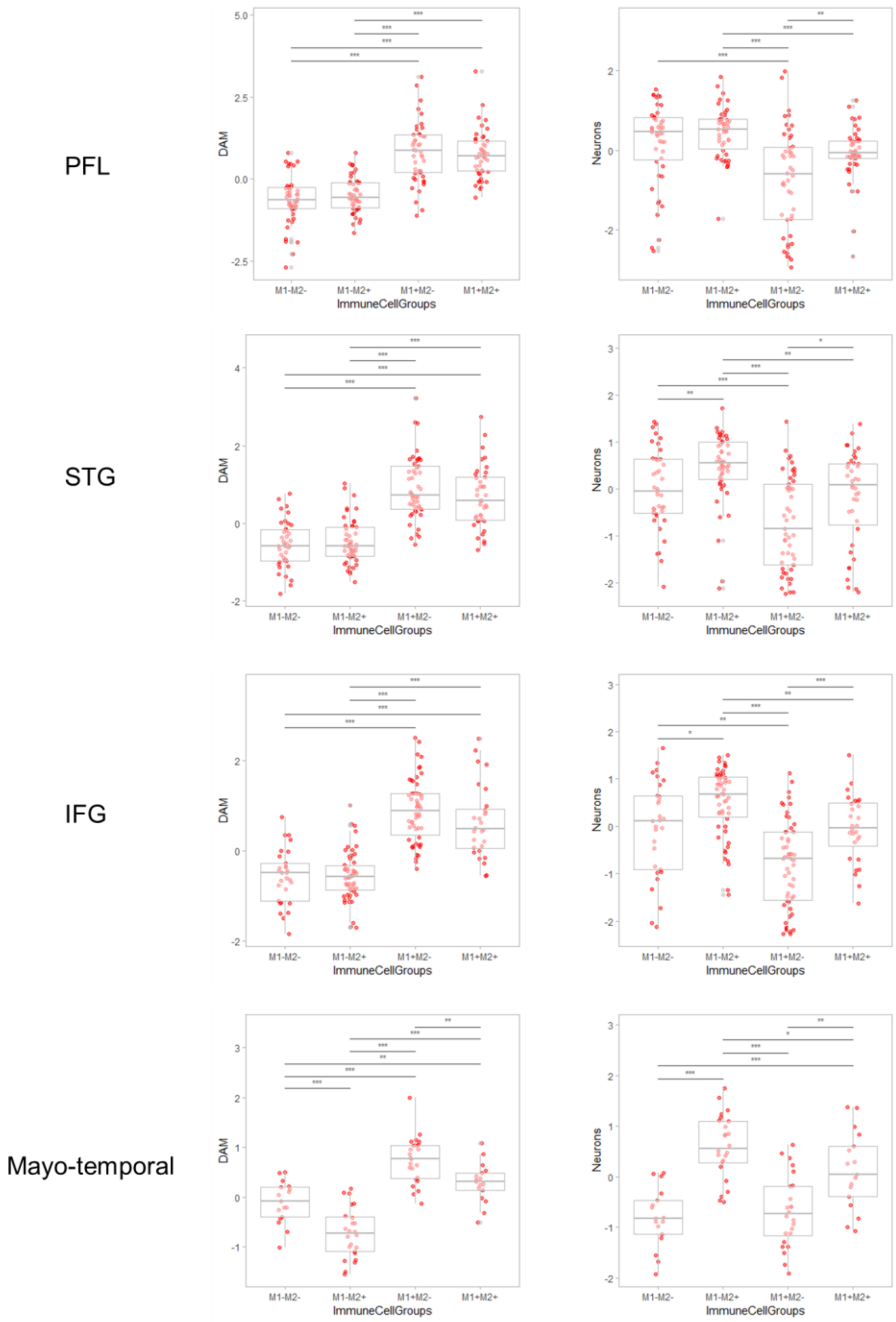


**Figure S4. Subgroups divided by M1- and M2-like cell enrichment.** The AD patients were divided into 4 subgroups according to the median value of M1- and M2-like cell enrichment scores. In all brain regions including PFC, STG, IFG and temporal cortex of Mayo RNA-seq data, DAM and neuron signatures were significantly different between the subgroups (F =50.1, $p < 1\times10^{-15}$ and F = 13.5, $p < 1\times10^{-9}$ for DAM and neuron, respectively in PHL; F =

51.6, $p < 1\times10^{-15}$ and $F = 14.2$, $p < 1\times10^{-9}$ for STG; $F= 60.5$, $p < 1\times10^{-15}$ and $F = 21.4$, $p < 1\times10^{-10}$ for IFG; $F = 44.3$, $p < 1\times10^{-15}$; $F = 21.0$, $p < 1\times10^{-9}$ for temporal cortex of Mayo data). Post-hoc analyses revealed that the M1+M2- subgroup showed the highest DAM enrichment and the lowest neuron signature in all brain regions (* for $p < 0.05$, ** for $p < 0.01$, and *** for $p < 0.001$; estimated by Post-hoc Tukey test).

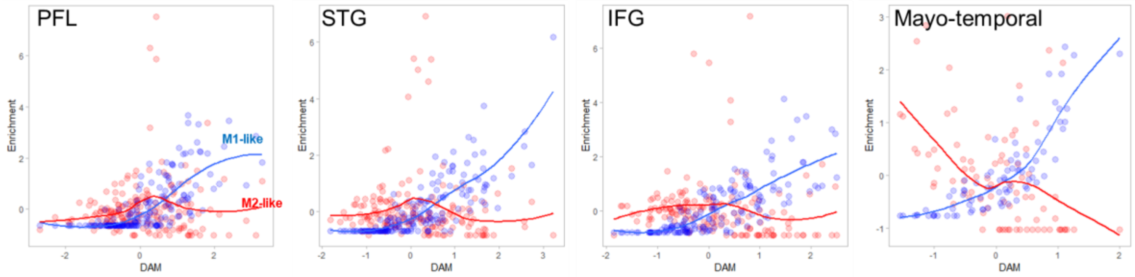


**Figure S5. M1- and M2-like cell signatures according to DAM enrichment in various brain regions.** M1- and M2-like cell enrichment scores were plotted with DAM scores in each brain region. M1-like cell was positively associated with DAM, while M2-like cell was not. M2-like cells were increased initially according to increased DAM, and then relatively decreased when DAM was further increased. M2-like cells showed negative correlation in Mayo temporal cortex data.

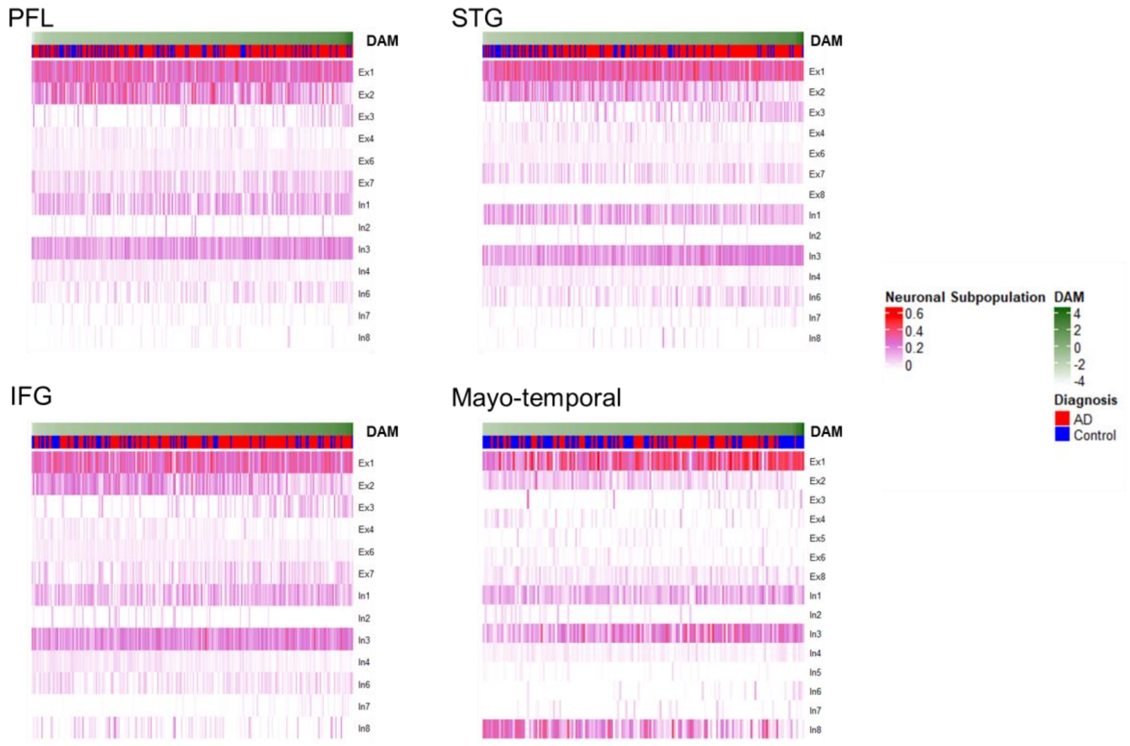


**Figure S6. The proportion of neuronal subpopulation according to DAM enrichment in various brain regions.** Neuronal subpopulation was digitally dissected by using genetic profiles of 8 excitatory and 8 inhibitory neuronal subtypes. For each brain region, the proportion of neuronal subpopulation was represented by a heatmap.

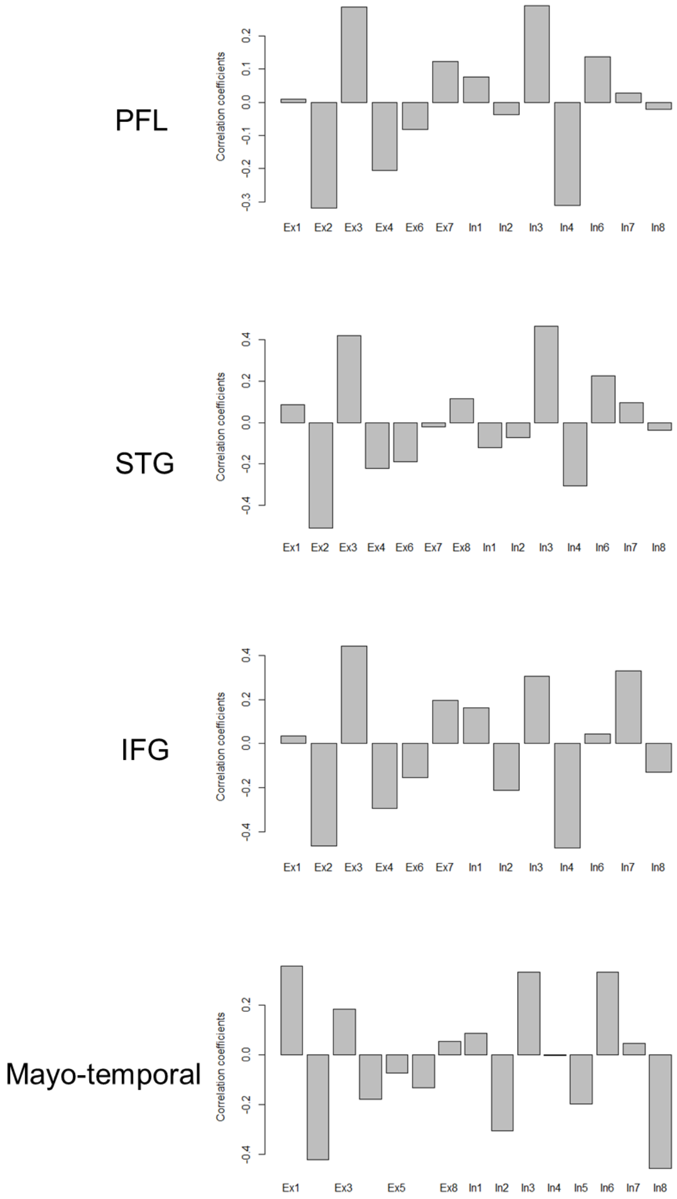


**Figure S7. Correlation coefficients between the proportion of neuronal subpopulation and DAM enrichment.** Though the correlation patterns of neuronal subpopulation were slightly different according to the brain regions, there was a common trend. The Ex3 proportion was positively correlated with DAM while the Ex2 proportion was negatively correlated with DAM in all brain regions.

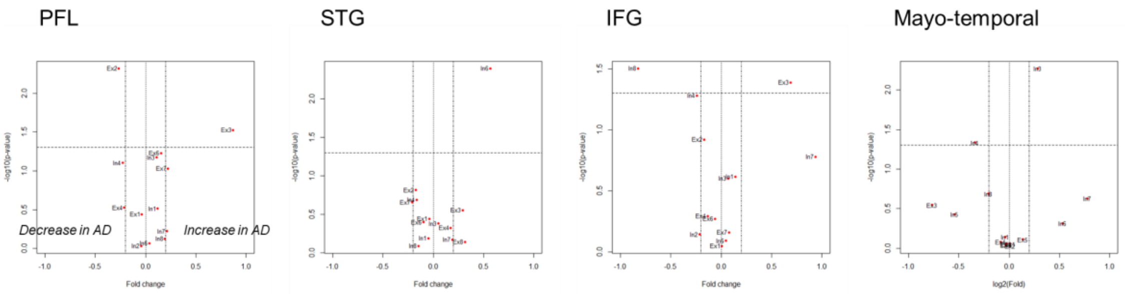


**Figure S8. Volcano plots for the comparison of the proportion of neuronal subtypes of AD and controls.** In AD brain, there was a trend of higher proportion of Ex3 neurons and lower proportion of Ex2 neurons. This pattern was found in PFL as well as PHG (Figure 4). However, in other brain regions including STG, IFG and temporal cortex, the proportion of these two excitatory neurons was not associated with the diagnosis of AD.
